# Proteomic Profiling of Rabid Canine Brain Samples Reveals Dysregulation of Immune Signalling and Neuronal Pathways

**DOI:** 10.64898/2025.12.02.691792

**Authors:** Ankeet Kumar, Sudeep Nagaraj, Baldev Raj Gulati, Gundallahalli Bayyappa Manjunatha Reddy, Utpal Tatu

## Abstract

Rabies is one of the most feared diseases and has been known to humans for approximately 4,000 years. It is caused by several lyssaviruses and leads to an encephalitis, which is 100% fatal after symptom onset. The most common cause of rabies is the rabies virus (RABV), which leads to around 59,000 human deaths globally every year. The infection leads to drastic changes in host behaviour, but the underlying mechanistic details remain unclear. Although interactions between the viral glycoprotein and host nicotinic acetylcholine receptors have been proposed as drivers of rabies-associated neurological dysfunction, clinical proteomic studies in naturally infected canines have been partially explored. In this study, we conducted a global proteomics analysis of RABV-infected and non-infected canine brain samples in India to unravel the changes in protein expression. Using liquid chromatography combined with mass spectrometry, we identified various host proteins and pathways dysregulated in the infected state. Approximately 700 proteins exhibited differential expression, with around 250 proteins involved in various pathways being significantly upregulated and approximately 500 proteins being significantly downregulated in the infected condition. Network clustering of the dysregulated proteins revealed functional modules, including clathrin-mediated endocytosis, actin cytoskeletal regulation, and the TCA cycle, indicating widespread alterations in cellular trafficking, energy metabolism, and structural integrity. Complementary pathway enrichment through PANTHER identified processes linked to axon guidance, integrin signalling, chemokine and cytokine-mediated inflammation, and cytoskeletal regulation, underscoring the profound impact of rabies virus infection on neuronal connectivity and host immune responses.

## Introduction

Rabies is a zoonotic viral encephalitis, which shows a 100% mortality rate after symptom onset. There is no treatment available for rabies, and the only preventive measure available for the disease is vaccines as pre-exposure prophylaxis (PrEP) and vaccines and rabies immunoglobulins (RIGs) as post-exposure prophylaxis (PEP). Rabies virus (RABV) belongs to a group of viruses in the order Mononegavirales and genus *Lyssavirus*. These viruses are bullet-shaped, enveloped viruses with negative-sense single-stranded non-segmented RNA as their genomes, ranging from 10 to 15 kb in size. RABV genome is approximately 12 kilobases (kb), coding for five proteins: nucleoprotein (N), phosphoprotein (P), matrix protein (M), glycoprotein (G) and large protein (L). These proteins are multifunctional and are known to interact with various host proteins [1]. Nucleoprotein enwraps the genome and is present as a multimer inside the virus [2]; it also associates with Heat-shock protein (Hsp), such as Hsp70, which is known to positively regulate viral infection and has been reported in viral nucleocapsids [3]. Phosphoprotein is a 297-amino-acid protein that acts as a cofactor for L protein and also binds to viral N protein to keep it soluble [4,5]. It also associates with host proteins like STAT and microtubules [6,7]. M protein binds to G and N protein and is deemed to be essential for the assembly and the bullet-shaped structure of the virus [8]. It is also known to interact with host histone deacetylase 6 [9]. G protein is present as a trimer on the virus and is a type I transmembrane protein; it can change between two forms influenced by pH change: closed and extended [10]. The G protein is known to bind to various host-receptors; major ones are N-Acetylcholine receptor, NCAM, p75NTR and Integrin [11-13]. Not all of these receptors are essential for RABV entry [11], and a recent study reports that RABV shows a preference for different host-entry receptors in the different hosts [14]. G protein is also the main protein against which the immune response is directed [15,16].

The virus is known to drastically alter the host’s behaviour and physiology [17], making it essential to investigate the host proteome under diseased conditions to understand the molecular mechanisms driving pathogenesis. In this study, we analysed brain samples from rabies virus-infected and non-infected dogs (*Canis familiaris*) from India. Comparative proteomic profiling revealed differential expression of over 750 host proteins. Enrichment analysis revealed significant alterations in pathways associated with endocytosis, synaptic signalling, neurotransmitter release, cytoskeletal regulation, antigen presentation, and cellular metabolism, including disruptions to the tricarboxylic acid (TCA) cycle and the respiratory electron transport chain. Additionally, proteins involved in angiogenesis, axon guidance, neurodegeneration-related processes, integrin signalling, and inflammatory signalling were markedly affected. These disruptions collectively suggest that rabies infection causes a collapse of neuronal communication, hijacks cellular metabolism, and initiates strong antiviral responses.

## Methods

### Sample preparation

The samples were tested for two genes, N and P, at the time of testing for the disease. These samples were transported to the lab and then tested for the N gene with a different set of primers. We used two brain samples (cerebrum) for each condition (infected vs. non-infected) in this study. Around 3g of brain sample was used for protein extraction using the Trizol, following the manufacturer’s protocol. Two technical replicates were processed for each sample by firstly chopping down the animal tissue into fine pieces. The RNA was stored for genomics, and the protein fraction was processed using the acetone precipitation method to isolate the total proteins in the sample. The total protein concentration was measured using the Bradford assay before proceeding with the proteomics.

### Trypsin digestion

Around 100 μg of protein was used for sample processing for each replicate. The protein was precipitated using acetone, followed by incubation with 10 mM Dithiothreitol (DTT) at 56°C for 45 minutes in a heat block with intermittent shaking. This was followed by 50 mM iodoacetamide (IAA) treatment at 37 °C for 30 minutes. The protein was then subjected to overnight trypsin digestion at 37 °C and 300 rpm (rotation per minute), at a 1:50 ratio (1 µg of trypsin per 50 µg of protein, using a 1 mg/mL stock), as per the protocol. The reaction is stopped the following day using formic acid.

### Desalting and peptide extraction

The resin columns are first activated using acetonitrile. Firstly, 150 μL of 50% Acetonitrile (ACN) is added and incubated for 2 minutes at room temperature. This step is done thrice, followed by equilibrating of the column with ACN and Trifluoroacetic acid (TFA) solution, followed by binding of the sample to the column. This is again followed by elution of the peptides using 70% ACN. The peptides are then dried and subjected to Liquid Chromatography-Mass Spectrometry (LC-MS).

### Liquid chromatography and tandem mass spectrometry

Desalted and dried samples were normalised to 0.2 µg/µL, and 10 µL of each sample was injected into the LC-MS system. Peptides were analysed in LC-MS Orbitrap fusion (Thermo Scientific). They were separated on a C-18 in-house column (15cm length, 1.5µ particle size, internal diameter 150µm). Peptides were eluted with a flow rate of 0.5µl/min, a gradient of 2%-90% ACN in 0.1% formic acid and analysed in positive-ion mode of the nano-electrospray ionisation source. The mass spectrometer was operated in a data-dependent mode, automatically switching between MS and MS/MS acquisition. Automated MS data of peptides were acquired through an Orbitrap mass analyser between 350 m/z and 2000 m/z above the 5,000 count threshold. The 20 most intense ions were selected for MS/MS acquisition using an ion trap mass analyser. The ion charge states were +2 to +8. The higher-energy C-trap dissociation (HCD) fragmentation energy was adjusted to 30%.

### Data analysis

The data analysis was performed using Proteome Discoverer v2.5, using the dog proteome (**UP000805418**) and RABV proteome accessed from UniProt on 05-04-2025. The viral databases used in this study are presented as **Supplementary File 1**. The samples were first checked for RABV proteins and then analysed separately against the dog proteome. The protein hits were then analysed for normalisation, statistical analysis, and visualisation using the web-based tool MetaboAnalyst v6.0 (https://www.metaboanalyst.ca/) [18] to check the differential protein expression in the host. The pathway enrichment analysis was performed by Panther (https://pantherdb.org/) [19], Reactome (https://reactome.org/) [20], gProfiler (https://biit.cs.ut.ee/gprofiler/gost) [21], Metascape (https://metascape.org/gp/index.html) [22], and STRING (https://string-db.org/) [23].

## Results

### Peptides for viral glycoprotein and phosphoprotein are abundant in the infected brain

Protein sequences retrieved from UniProt were used as the reference database for peptide identification. Mass spectrometry analysis of infected samples consistently detected multiple viral proteins across biological and technical replicates, including the nucleoprotein, phosphoprotein, glycoprotein, and the large (L) protein. Among these, the nucleoprotein, glycoprotein and phosphoprotein were the most frequently and abundantly detected across all replicates, indicating their high expression or stability during infection. Notably, the large (L) protein and glycoprotein were also detected in several replicates, although with more variability. **Figure 1 A-D** shows the viral protein peptides found for the samples.

**Figure 1.**
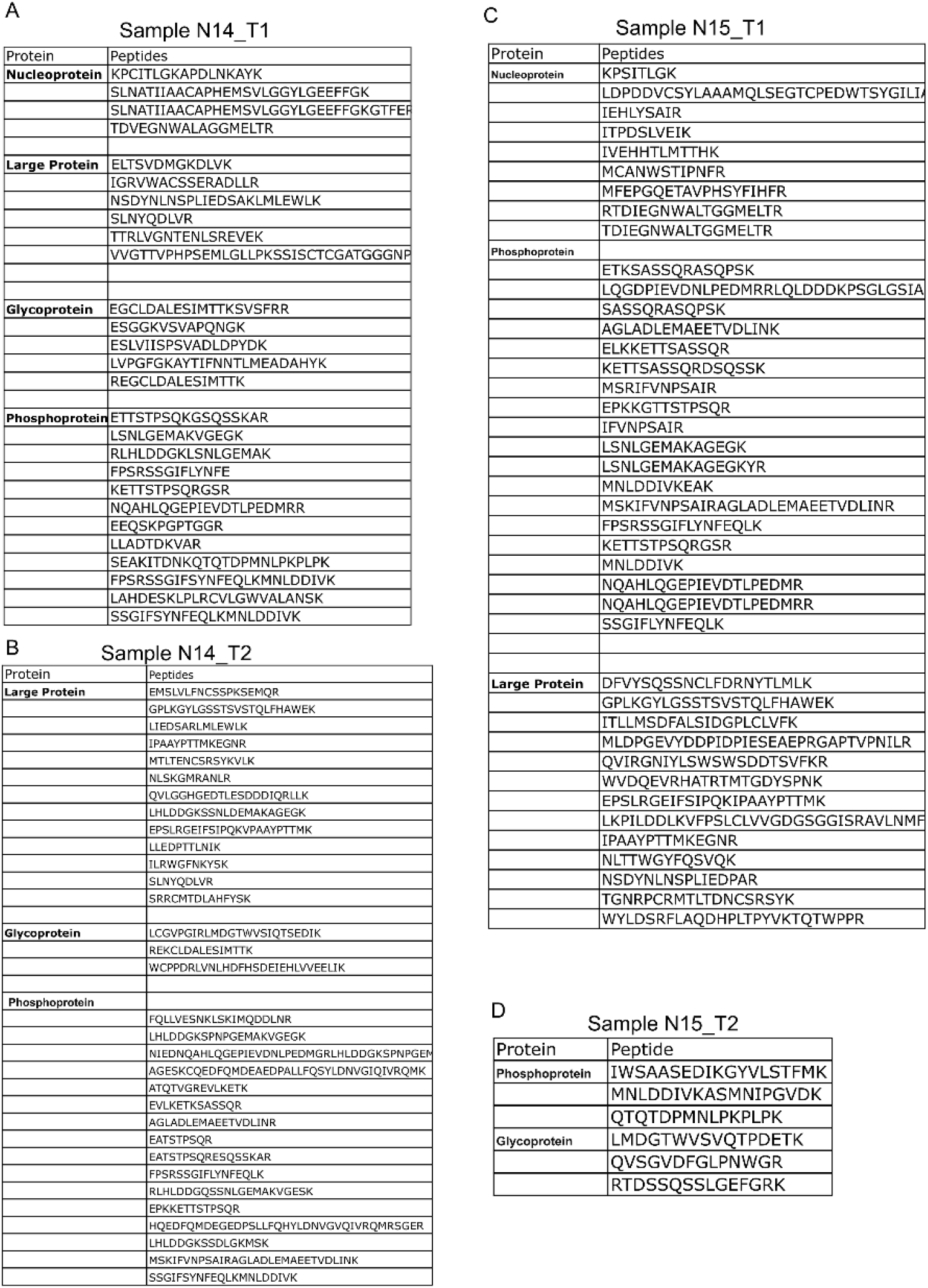
The peptides corresponding to viral proteins were detected in the infected canine samples. The criteria used for protein detection were more than 3 unique peptides per protein. The sequence for the peptides identified for viral proteins is presented in the tables for each sample

### Around 750 host proteins were found to be differentially expressed in the rabid brain samples

Fold change analysis of protein expression intensities identified around 250 significantly upregulated and around 500 significantly downregulated proteins in infected samples compared to non-infected controls. The complete list of differentially expressed proteins is provided in **Supplementary File 2**. To visualise the distribution and significance of protein expression changes, a volcano plot (**Figure 2A**) was generated. The plot highlights the ten most differentially expressed proteins with statistically significant p-values. Among the upregulated proteins, BRCA-associated protein, outer dense fiber protein, and Hippocalcin showed the highest fold changes. Conversely, Aminopeptidase, Sec7 domain-containing protein, and Wnt signalling-related protein were among the most downregulated candidates.

**Figure 2.**
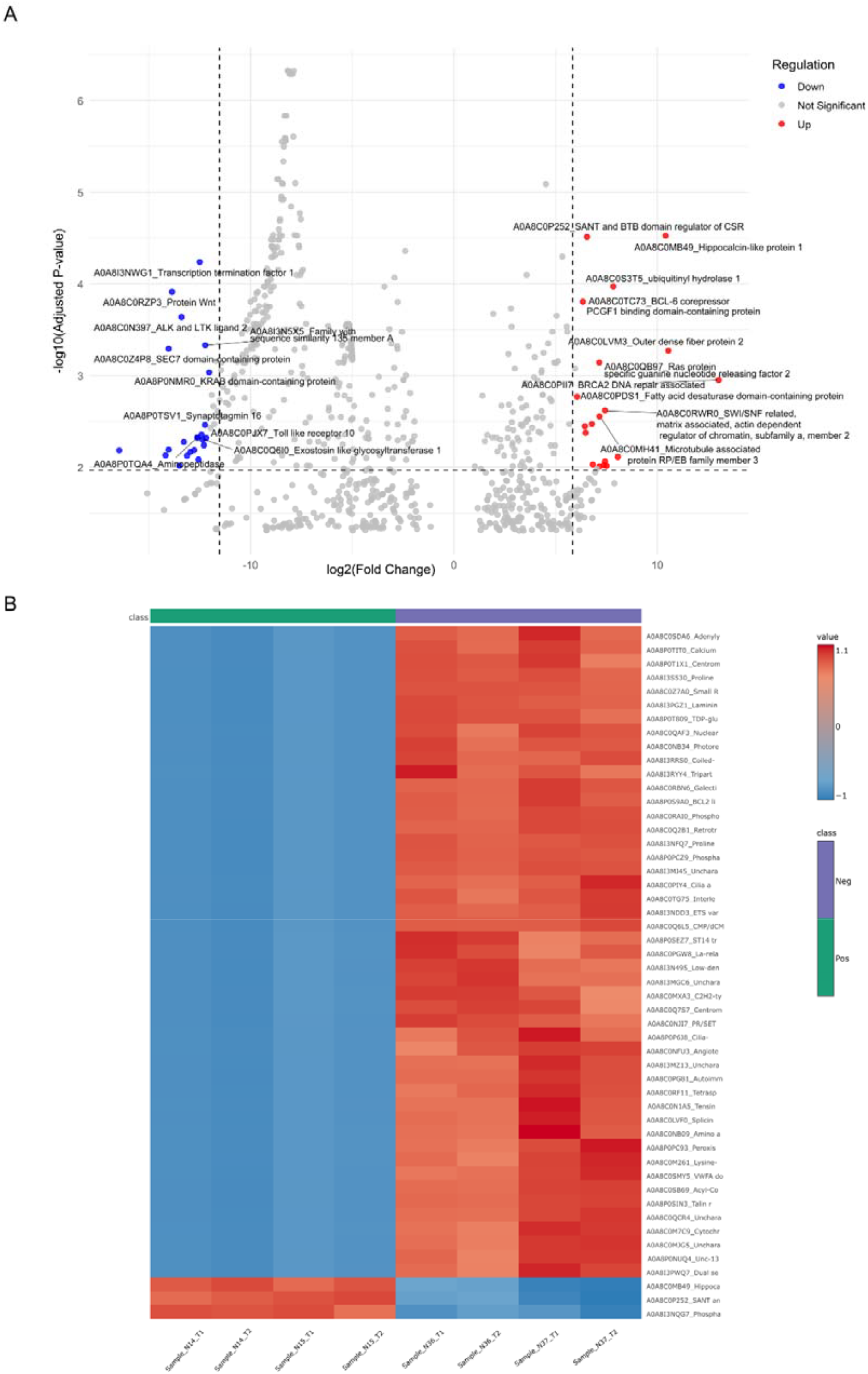
Differentially expressed proteins in infected canine samples. **(A)** Volcano plot showing significantly upregulated (red), downregulated (blue), and non-significant (grey) proteins based on thresholds of -log□□(adj. p-value) > 2 and log□ fold change (log□FC) > 6 for upregulation, and log□FC < -12 for downregulation. **(B)** Heatmap displaying the expression profiles of the top 50 differentially expressed proteins in infected versus non-infected samples.

**Figure 2B** presents a heatmap of the top 50 differentially expressed proteins, illustrating clear separation between infected and non-infected samples. Notably, several of the top upregulated proteins in infected samples are involved in neural signalling, cytoskeletal organisation, and pathways regulating chromatin remodelling and transcriptional control. **Table 1** summarises proteins showing concordant expression trends in previous studies and our current study.

**Table 1.** Comparison of dysregulated host proteins identified in our study with previous reports.

| Serial Number | Protein | Expression | Publication |
| --- | --- | --- | --- |
| 1 | Apolipoprotein E | Downregulated | Chienwichai P, 2025 [24] |
| 2 | Hsp90 and Hsp70 | Upregulated | Thanomsridetchai N, 2011 [25] |
| 3 | Dihydrolipoyl dehydrogenase | Upregulated | Chienwichai P, 2025 |
| 4 | Annexin | Upregulated | Chienwichai P, 2025 |
| 5 | Glutathione S-transferase | Upregulated | Thanomsridetchai N, 2011 |
| 6 | Peroxiredoxin-like 2A | Downregulated | Behera S, 2022 [26] |
| 7 | Malic enzyme | Upregulated | Venugopal A, 2013 [27] |
| 8 | Histone H4 | Upregulated | Venugopal A, 2013 |
| 9 | protein kinase C | Downregulated | Venugopal A, 2013 |
| 10 | Eukaryotic translation initiation factor 4 gamma 1 | Upregulated | Venugopal A, 2013 |
| 11 | Tetraspanin | Downregulated | Behera S, 2020 [28] |
| 12 | 4-aminobutyrate aminotransferase | Upregulated | Behera S, 2020 |

### Enrichment analysis reveals dysregulation of chemokine signalling and axon guidance pathways

Various host pathway-related proteins were seen to be dysregulated. **Figure S1** illustrates the protein classes that are upregulated and downregulated in the infected condition. **Figure 3A** shows a Venn diagram illustrating the number of pathways upregulated and downregulated in the RABV-infected samples, as classified by Panther. A total of twenty-two pathways were found for upregulated proteins, and twenty-six for downregulated proteins. Eight pathways were commonly affected in both the conditions (Angiogenesis, Integrin signalling pathway, Inflammation mediated by chemokine and cytokine signalling pathway, EGF receptor signalling pathway, Cytoskeletal regulation by Rho GTPase, Cadherin signalling pathway, Huntington disease, Wnt signalling pathway). Uniquely upregulated pathways belonged to immune-related, cytoskeletal system-related and cell signalling-related pathways, like apoptosis, toll-like receptors, and were downregulated in the infected condition, while several pathways involving neuronal signalling like nicotinic acetylcholine receptor signalling and oxytocin receptor mediated signalling pathways were only found for downregulated proteins.

**Figure 3.**
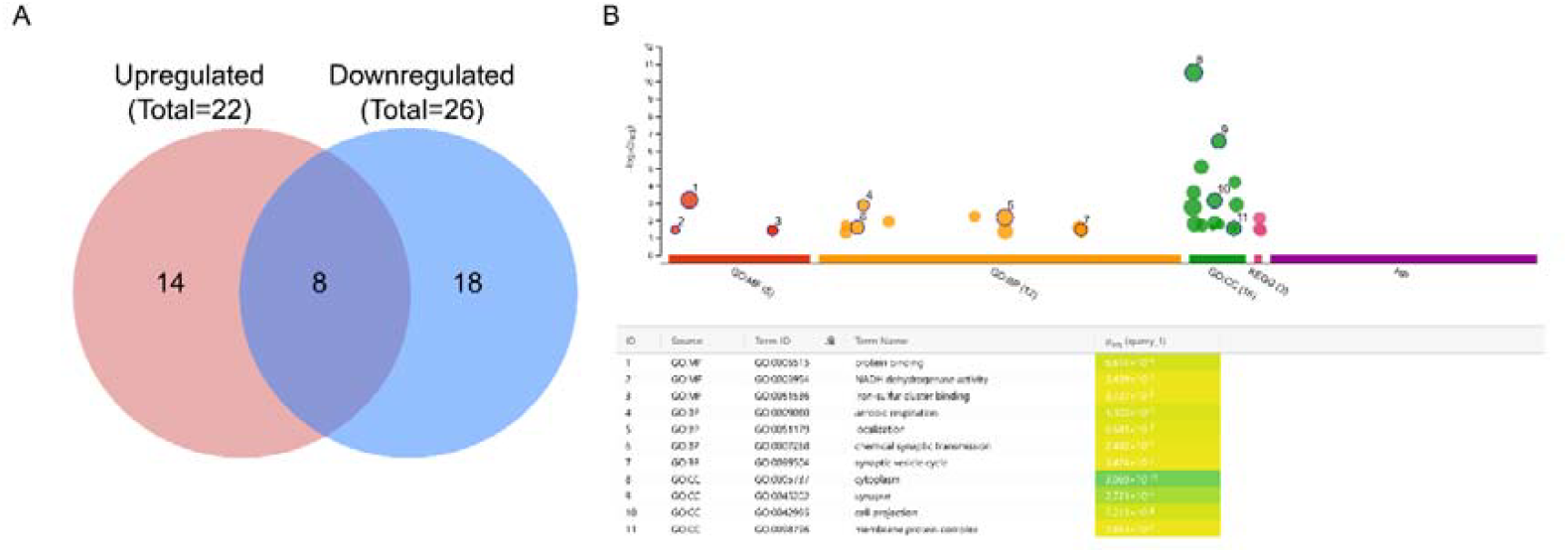
Pathway enrichment for rabid canine brain. **(A)** The Venn diagram shows the total pathways hit given by the proteins present in our dataset and their distribution across the two categories in the infected samples in Panther. (B) The upregulated protein pathways using gprofiler.

**Figure 3B** illustrates the enrichment of pathways and cellular components that are significantly upregulated in the infected condition. Among the molecular functions, proteins associated with protein binding, NADH dehydrogenase activity, and iron-sulfur cluster binding were enriched. The biological processes include aerobic respiration, localisation, and synaptic transmission. Notably, cellular components such as cytoplasm, synapse, cell projection, and membrane protein complex show strong enrichment. These findings collectively point to the upregulation of metabolic pathways, synaptic function, and structural cell components under infection.

### Network-level insights into host proteome dysregulation of endocytosis-related and antigen presentation pathways

We constructed a protein-protein interaction (PPI) network using STRING for the dysregulated host proteins identified in rabies-infected canine brain samples. The network comprised 280 nodes and 376 edges, with an average node degree of 2.69 and an average local clustering coefficient of 0.366. Notably, the number of observed edges (376) was markedly higher than the number of edges expected by chance (267), yielding a highly significant PPI enrichment p-value of 2.02 × 10□^1^□. This enrichment suggests that the dysregulated proteins are not randomly distributed but are functionally connected, indicating a coordinated involvement in shared or related biological pathways.

Figure 4. shows 37 gene expression clusters, each defined by a unique set of co-expressed genes and enriched biological functions. Several clusters were mapped to well-defined cellular processes, including aminoacyl-tRNA biosynthesis and translation (Cluster 3), the citric acid cycle and electron transport (Cluster 4), actin binding (Cluster 6), and antigen processing and presentation (Cluster 8). Other clusters corresponded to endocytosis (Cluster 2), spliceosome regulation (Cluster 9), RIG-I-like signalling (Cluster 11), ECM-receptor interaction (Cluster 12), and Hippo signalling (Cluster 26), among others. Notably, Cluster 1 was gene-rich (18 genes), including several related to centrosomal function and intracellular transport (e.g., CENPF, KIF9, IFT81), while some clusters (e.g., Clusters 21, 22, 27, 29) were more ambiguous or comprised less-characterised genes and functions. Interestingly, a few clusters (Clusters 16 and 25) indicated poorly understood or mixed functional annotations, suggesting the presence of novel regulatory modules. These findings provide a systems-level snapshot of the transcriptional landscape, highlighting both canonical pathways and less explored gene networks. A detailed description of clusters in a decreasing order of protein hits is presented in **Supplementary File 3**.

**Figure 4.**
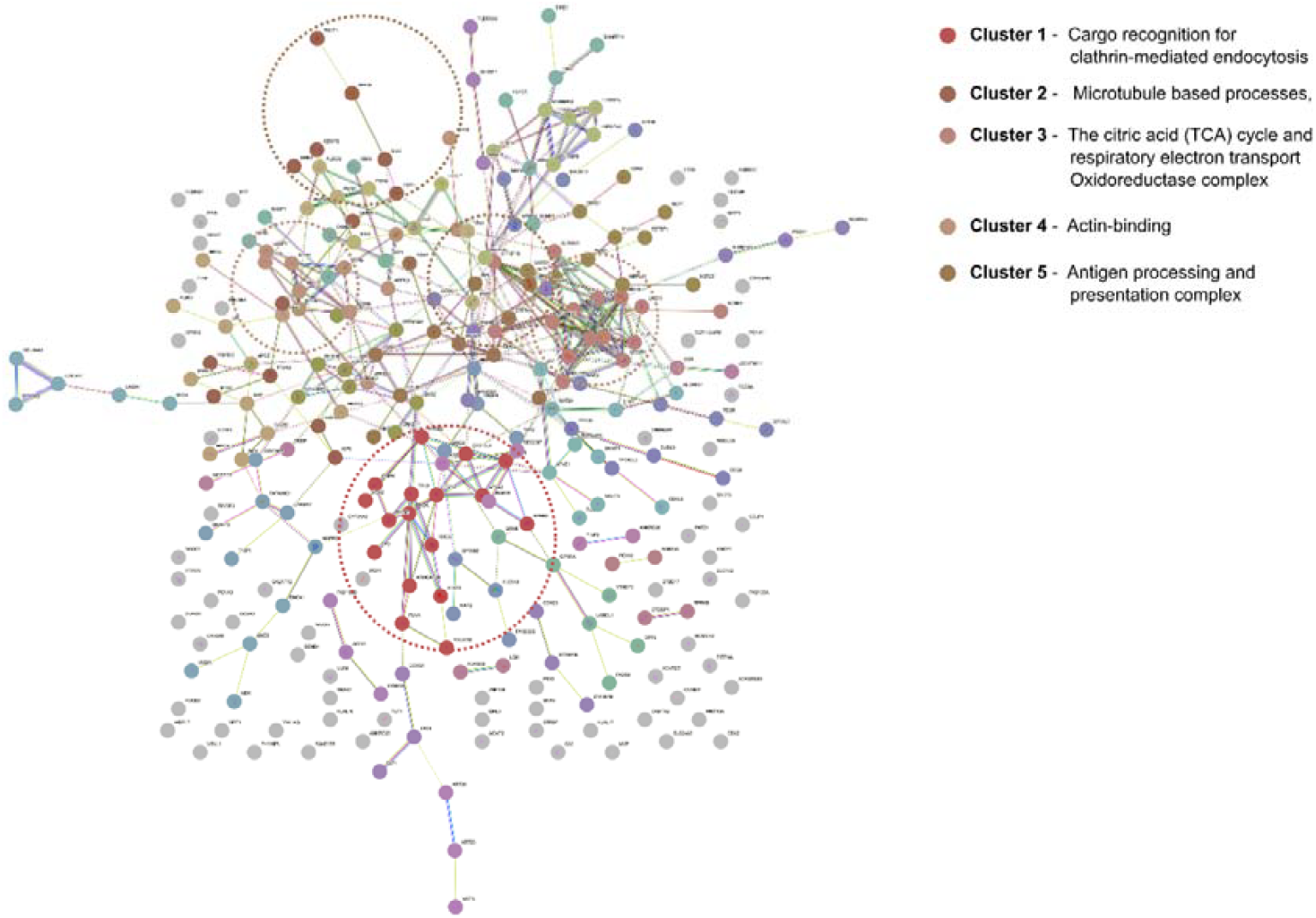
The protein-protein interaction network for dysregulated proteins in rabies. The network shows interactions among differentially expressed host proteins identified in infected samples. Clustering was performed using the Markov Clustering (MCL) algorithm with an inflation parameter of 1.5, which groups proteins into natural clusters based on their connectivity. Each cluster represents a group of proteins with potentially shared or related biological functions. Edges between nodes represent known or predicted protein-protein interactions derived from STRING data.

## Discussion

Despite being one of the oldest zoonotic infections known to mankind, mechanistic details related to symptoms and clinical pathogenesis are still lacking in rabies. Understanding these mechanisms requires a systems-level approach, and proteomics provides a powerful platform for capturing global alterations in host protein expression and pathway regulation during infection. In this study, we leveraged proteomic profiling of canine brain tissues to investigate host responses to RABV infection. By comparing infected and non-infected samples, we identified approximately 750 dysregulated host proteins, including known players such as Hsp70 and Hsp90, which are known to bind the viral nucleoprotein and support RABV replication [3,29]. Cross-referencing with previously published proteomic datasets is presented in **Table 1**.

In our dataset, we identified hits for several immune-related pathways, including the toll-like receptor pathway, chemokine signalling, and interleukin-associated proteins, which were found to be dysregulated. In addition to this, we found that a protein called Phosphoprotein Enriched in Astrocytes 15 (PEA-15) was also found to be ∼6-fold upregulated in RABV-infected brains. PEA-15 is a multifunctional scaffold protein with critical roles in regulating cell survival and proliferation [30]. It is also involved in cell growth regulations also has potent anti-apoptotic capabilities [30,31]. The dysregulation of these pathways and the upregulation of PEA-15 in RABV infection is particularly noteworthy given that RABV, especially street strains, are known to adopt a ‘stealth’ strategy characterised by the suppression of apoptosis during early infection [31,32]. The virus thus keeps the neurons alive in the host so that it has enough time for replication and spreading, while also not attracting the immune system very early [33]. The regulation of the host immune system by the virus is crucial for infection to occur and transmit; thereby, a drastic change is observed in the host proteome pertaining to the immune system during viral infection.

A large subset of dysregulated proteins belonged to the cytoskeletal network, including subunits of the Arp2/3 complex, MAPRE3, dynein axonemal heavy chain 3, actinin alpha 2, and microtubule-actin crosslinking factor 1. These changes highlight extensive remodelling of actin and microtubule dynamics during infection. Such reorganisation is a well-established viral strategy to facilitate intracellular transport, replication, and virion assembly. Consistent with earlier reports [34,35], our findings suggest that RABV actively manipulates the cytoskeleton to drive early disease progression and neuronal dysfunction. Subsequent functional enrichment analyses revealed a complex and multifaceted host response. Pathways related to energy metabolism, including aerobic respiration and NADH dehydrogenase activity, were significantly upregulated, possibly reflecting increased energy demands imposed by the virus. Downregulation of synaptic signalling and neuronal membrane proteins, such as synaptotagmin-like 1 and synaptotagmin 16, highlights potential disruptions to neural function, correlating with the neurological symptoms observed in rabies. Synaptic machinery is increasingly recognised as a druggable target for rabies [24]. Additionally, downregulation of the Notch signalling pathway [36], known for regulating glial activation and neuropathic pain [37], suggests potential virus-mediated immunomodulation. The role of the Notch pathway in RABV infection is not well understood. Interestingly, the nicotinic acetylcholine signalling pathway is also dysregulated. The nicotinic acetylcholine receptor (nAChR), which was the first host receptor identified for RABV and a known viral target associated with the behavioural changes seen in rabies (such as aggression and hydrophobia) [11,38], was also affected in the infected state. Additionally, neurodegenerative disease-associated genes involved in pathways, such as those associated with Huntington’s and Alzheimer’s diseases, were also significantly altered, further implying virus-induced modulation of brain homeostasis. These pathways have already been reported in rabies proteomics studies [27,28].

Interestingly, several pathways were enriched in both upregulated and downregulated datasets, including integrin signalling, angiogenesis, and chemokine-mediated inflammation. This bidirectional regulation within single pathways may reflect the dynamic interplay of viral infection and host responses, emphasising the need to study temporal dynamics during infection. Additionally, the upregulation of axon guidance molecules during infection may reflect the virus’s exploitation of host signalling for cell entry and intracellular trafficking. This finding aligns with recent insights showing that RABV engages the NRP2-TGFBR1 axis to trigger actin polymerisation [39], promoting clathrin-mediated endocytosis [40]. The convergence of axon guidance and viral entry signalling illustrates how RABV hijacks neuronal pathways to facilitate infection.

PPI network analysis further supported the notion of virus-induced functional reprogramming. Dysregulated proteins clustered around key cellular systems, including clathrin-mediated endocytosis, actin filament organisation, heat shock-related, protein folding, RNA binding, antigen binding and processing, voltage-gated channels and splicing-related. Cargo recognition for clathrin-mediated endocytosis and extracellular matrix (ECM)-receptor interaction was particularly affected, a process essential for viral entry. Moreover, the involvement of dynein, a microtubule motor protein crucial for retrograde axonal transport, points to hijacking of intracellular trafficking machinery by the virus. Several immune-related hubs, such as the RIG-I-like receptor signalling pathway, were also disrupted, suggesting immune evasion or delayed antiviral activation. These network-wide disruptions illustrate how RABV systematically remodels host cellular infrastructure to create a pro-viral environment. Furthermore, RABV is known to form protective structures called Negri bodies, a diagnostic feature of infection, within which the virus replicates at high titers while evading the host immune response [4,5,41].

Our data also provide insights into lesser-known players in rabies infection. For instance, the SWI/SNF complex component SMARCA2 has known roles in other viral systems (e.g., HIV, RSV) but has not yet been explored in rabies [41-43]. Proteins involved in redox regulation and oxidative stress, such as SH3 domain-binding glutamic acid-rich-like protein, glutathione peroxidase, and glutathione reductase, were altered, suggesting that oxidative stress may be a significant aspect of rabies pathogenesis [45]. The downregulation of vimentin, an important mediator of viral attachment and immune signalling [45-47] and annexin A2 further supports alterations in host structural and immune systems. In addition, ubiquitin-related proteins such as UFM1 and UBA6 were upregulated, consistent with the role of ubiquitin signalling in viral infections[48-50].

In conclusion, the proteomic landscape of RABV infection in the canine brain reveals widespread remodelling of host cellular systems critical for neuronal function and survival. Key pathways, such as metabolism, cytoskeletal organisation, synaptic signalling, and immune regulation, have been significantly disrupted. The virus appears to exploit core host processes, including clathrin-mediated endocytosis, axonal transport, and immune evasion, to facilitate neural invasion and persistence. Notably, synaptic communication and structural integrity were prominently suppressed in infected brains, with metabolic and immune dysfunction, which all together likely underpin rabies-induced neuropathology. The findings establish new directions for identifying host-derived biomarkers and therapeutic targets and emphasise the need for further mechanistic studies of the newly implicated host factors driving rabies pathogenesis.

## Acknowledgements

The authors thank Utpal Tatu lab members for their discussions and input.

## Author contributions

Conceptualisation and Study Design: Ankeet Kumar, Gundallahalli Bayyappa Manjunatha Reddy and Utpal Tatu

Experimental Design, Facility Generation: Baldev Raj Gulati, Utpal Tatu

Data curation: Ankeet Kumar and Gundallahalli Bayyappa Manjunatha Reddy

Sample collection, Testing and Archiving: Sudeep Nagaraj and Gundallahalli Bayyappa Manjunatha Reddy

Formal analysis: Ankeet Kumar and Gundallahalli Bayyappa Manjunatha Reddy

Investigation: Ankeet Kumar, Gundallahalli Bayyappa Manjunatha Reddy and Utpal Tatu

Methodology: Ankeet Kumar and Utpal Tatu

Supervision: Utpal Tatu

Project administration: Utpal Tatu

Resources: Utpal Tatu, Gundallahalli Bayyappa Manjunatha Reddy

Validation: Ankeet Kumar, Gundallahalli Bayyappa Manjunatha Reddy and Utpal Tatu

Writing - original draft: Ankeet Kumar

Writing - review & editing: Ankeet Kumar, Gundallahalli Bayyappa Manjunatha Reddy and Utpal Tatu

## Data Availability

The data supporting the findings in the study are available in the manuscript and as supporting data. The raw data will be made available upon request.

## Ethics approval

Permissions for animal sampling were granted by the concerned authorities, the Indian Institute of Science, under approval number CAF/Ethics/831/2021 and NIVEDI, under the approval number NIVEDI/IAEC/2025/15.

